# Eating to dare - Nutrition impacts human risky decision and related brain function

**DOI:** 10.1101/2020.05.10.085688

**Authors:** Lu Liu, Sergio Oroz Artigas, Anja Ulrich, Jeremy Tardu, Peter N. C. Mohr, Britta Wilms, Berthold Koletzko, Sebastian M. Schmid, Soyoung Q. Park

## Abstract

Macronutrient composition modulates plasma amino acids that are precursors of neurotransmitters and can impact brain function and decisions. Neurotransmitter serotonin has been shown to regulate not only food intake, but also economic decisions. We investigated whether an acute nutrition-manipulation inducing plasma tryptophan fluctuation affects brain function, thereby affecting risky decisions. Breakfasts differing in carbohydrate/protein ratios were offered to test changes in risky decision making while metabolic and neural dynamics were tracked. We identified that a high-carbohydrate/protein meal increased plasma tryptophan which mapped to individual risk propensity changes. Moreover, the meal-driven fluctuation in tryptophan and risk propensity changes were modulated by individual difference in body fat mass. Using fMRI, we further identified activation in the parietal lobule during risk-processing, of which activities 1) were correlated with the risk propensity changes in decision making, 2) were sensitive to the tryptophan fluctuation, and 3) were modulated by individual’s body fat mass. Furthermore, the activity in the parietal lobule positively mediated the tryptophan-fluctuation to risk-propensity-changes relationship. Our results provide evidence for a personalized nutrition-driven modulation on human risky decisions and its metabolic and neural mechanisms.

## Introduction

Food intake profoundly impacts not only human physiological processes but can also change our behaviors. The macronutrient composition of a food modulates multiple biochemical processes, among others plasma concentrations of large neutral amino acids (LNAAs) that can pass the blood-brain barrier. Specifically, tryptophan is the precursor of neurotransmitter serotonin (5-hydroxytryptamine, 5-HT) (1). Across species, plasma tryptophan has been shown to be closely related to the brain serotonin level (2, 3). Importantly, a meal with a higher carbohydrate/protein ratio increases plasma tryptophan level (1, 4), which presumably enhances brain serotonin level. A change in serotonin dynamics in the brain has been shown to impact specific brain functions and behaviors, ranging from food intake to aversive processing (5, 6). Although this provided novel possibilities for neuro- and behavioral modification through nutrition intervention, so far empirical evidence is lacking (7).

The central serotonergic system is targeted in major neuropsychiatric treatments and has been shown to modulate decision making (8, 9), but effects are mixed. By serotonergic manipulation, some studies indicated that serotonin facilitates punishment processing and behavioral inhibition (10, 11). In line with this, a pronounced risk, loss or harm avoidance was associated with enhancement of central serotonin (5, 12, 13). However, other studies showed that serotonin increased risk-seeking or loss-chasing after manipulation of serotonergic systems (14, 15). Considering the complex roles of serotonin in energy homeostasis, a potential explanation is that serotonergic modulation on risk or harm aversion processing might be complex. Supporting this, Crockett and colleagues (5) has reported the application of an acute dose of citalopram (a selective serotonin reuptake inhibitor, SSRI) changed harm aversion in body-weight dependent manner. In line with this, individual’s risk preference has been shown to depend on the physiological status, such as body energy reserve (16–18). Specifically, both the metabolic states (fasting vs satiety) and energy reserves influenced individual’s risk propensity during decision making (17, 18). Furthermore, energy homeostasis is implicated to be closely associated with serotonergic system (6, 19). Central serotonin inhibits food intake and increases energy expenditure (6), whereas peripheral serotonin may upregulate energy storage (20). For example, studies have reported positive correlation between peripheral serotonin levels and obesity/lipid levels in animal models (20, 21). Taken together, the decisions made in the brain may integrate internal state (i.e. energy reserve) and the environment (i.e. property of stimuli) (7). However, it is still unknown whether and how the nutrient affected fluctuation of serotonin precursor tryptophan relates to body composition and to risk decision performance.

Moreover, quite few studies focusing on the effects of serotonergic system on decision making has explored the underlying brain functions in healthy human participants (22–24). Evidence from previous studies about risky decision making have identified many brain regions including the frontal cortex, parietal cortex, insular cortex, and amygdala in reward risk processing; and regions including the ventromedial prefrontal cortex (vmPFC), orbital frontal cortex (OFC) and striatum in outcome evaluation (25–28). Specifically, studies with decision making tasks have suggested the possible involvement of dorsomedial prefrontal cortex (dmPFC), ventrolateral prefrontal cortex (vlPFC) and parietal cortex in risk preferences processing (25, 27–29). In line with this, a rat study manipulating the serotonergic system in the insular and OFC identified risk preferences changes during decision making (30). However, whether and how the impact of nutrient-composition on serotonin precursor tryptophan relates to brain function changes and to decision performance is still lack of investigation.

In this study, we experimentally manipulated the macronutrient composition of a standardized breakfast in a double-blind within-subject study design and tested whether a one-shot meal intervention can modulate risk propensity. Importantly, we systematically merge the metabolic, neural and behavioral data to dynamically capture the nutrition-caused changes and their dynamics. To do so, we take into account the temporal dynamics of plasma tryptophan and body fat mass. Furthermore, the changes in neural processing during the decision making as a function of macronutrient manipulation were captured by means of functional magnetic resonance imaging (fMRI). In line with serotonergic pharmaco-intervention studies showing that 5-HT modulated risky decisions, we hypothesized that the high-carb/protein ratio breakfast would also change the risk propensity, which attribute to the plasma tryptophan fluctuation. Furthermore, we tested possible effects of body fat on the macronutrient-driven tryptophan fluctuation, as well as the risk behavior changes. On neural level, we aimed to unveil brain regions, in which activity reflects the nutrient-driven changes in metabolism and behavior. To enable this, we searched for brain regions of which activity fulfills all three criteria: The activity of the brain region 1) codes the nutrition-affected risk behavior changes, 2) tracts the dynamics in plasma tryptophan and 3) reflects the individual differences in body fat.

## Results

### The Influence of Macronutrient Composition on Plasma Parameters

First, we checked whether the different meals induce different changes in tryptophan temporal dynamics: large neutral amino acid (Tryp/LNAA) ratio, by computing a repeated-measures ANOVA with time (T1-T4) and meal session (high vs. low-carb/protein breakfasts) as within-subject factors. We observed a significant increase in plasma tryptophan after a high-carb/protein breakfast. Specifically, there was a main effect of condition [*F*(1, 34) = 66.23, *p* < 0.001], a main effect of time [*F*(3, 102) = 10.99, *p* < 0.001] and an interaction between condition and time [*F*(3, 102) = 16.52, *p* < 0.001] in tryptophan/LNAA ratios. Post hoc analysis revealed higher tryptophan levels after the high-carb/protein breakfast for all postprandial samples [T2: *t*(34) = 7.24, *p* < 0.001; T3: *t*(34) = 8.32, *p* < 0.001; T4: *t*(34) = 8.06, *p* < 0.001] (Fig. 2*A*). We also observed in a repeated-measures ANOVA that the individual time course of blood glucose was significantly modulated by the respective macronutrient condition [session X time interaction; *F*(7, 238) = 7.39, *p* < 0.001] (Fig. 2*B*). Contrary to tryptophan, the tyrosine increased with greater protein content of the meal (see Results section in *SI Appendix*). For the other metabolic parameters [insulin, cortisol, ghrelin, and leptin], there were no significant modulation by session (*SI Appendix*, Table S1).

**Fig. 1.**
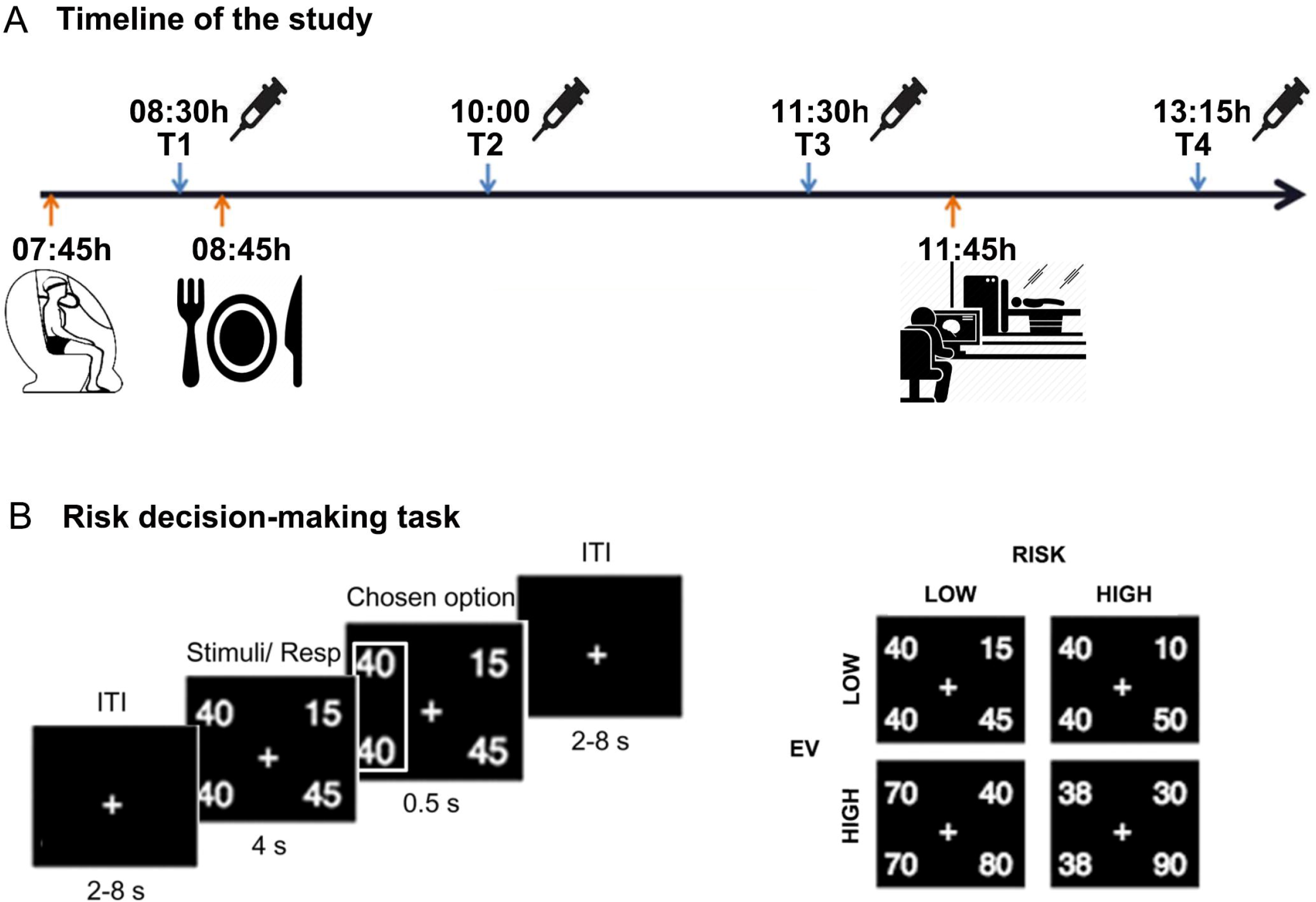
Study design. (A) Timeline of the study. Subjects attended the research unit at 0730 hours and were prepared for the experiments. At 0845 hours subjects received breakfast according to the respective experimental condition. Blood was drawn at 8 time points. T1–T4 indicates blood samples used for measurement of tryptophan and tyrosine. (B) The risk decision making task procedure.

**Fig. 2.**
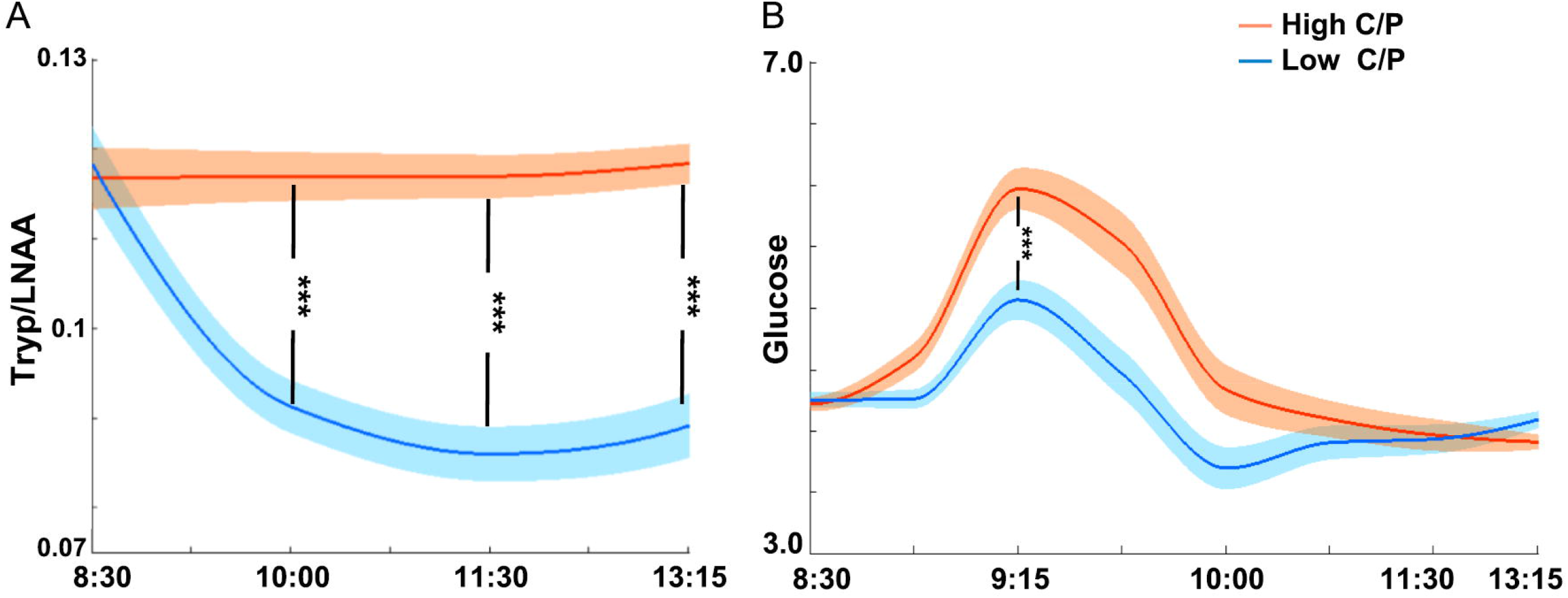
Nutrient manipulation on postprandial tryptophan/LNAA and glucose levels. Macronutrient composition-dependent changes in postprandial tryptophan/LNAA and glucose (mmol/L) levels. Red lines indicate high-carb/protein session and blue lines indicate low-carb/protein session for (A) tryptophan/LNAA ratio; (B) glucose concentrations (±SEM in shadowed area). High C/P, high-carb/protein breakfast; Low C/P, low-carb/protein breakfast; Tryp/LNAA, tryptophan/LNAA ratio. ^*^*P* < 0.05, ^**^*P* < 0.01 and ^***^*P* < 0.001.

Next, we assessed the effect of body fat mass (mean ± s.d. = 13.78 ± 5.22 kg) on nutrient-driven tryptophan metabolism by using linear regression model with breakfast sessions and body fat mass as regressors. A significant session by fat mass interaction was confirmed (β = 0.38, SE = 0.13, *t* = 2.99, *P* = 0.004). That is, the fat mass modulated the relationship between macronutrient-manipulation and tryptophan/LNAA fluctuation (see *SI Appendix*). In addition, to assess the insulin resistance/sensitivity of the participants, a Homeostatic Model Assessment of Insulin Resistance (HOMA-IR) was conducted and we found the mean score of HOMA-IR was 0.79 (0.42 - 1.50), suggesting that the participants were insulin sensitive.

### The Effect of Macronutrient Composition on Risky Decision Making Modulated by Fat Mass

The rate of risk choices were analyzed using t-tests and ANOVAs. In each condition, participant’s risk propensity was defined as the proportion of risk-taking trials (29). The risk rates in a session by value level (low, high) by risk level (low, high) ANOVA was conducted, and no significant effects of session nor session by condition interactions were found, nor the session effect on risk aversion (*Materials and Methods* and *SI Appendix*, Table S2). According to previous literature (5, 18), we hypothesized that the individual difference in body fat mass would modulate the effect of macronutrient on risk behaviors. We first compared the mixed effects regression models (*Materials and Methods* and *SI Appendix*, Table S3) in explaining participants’ risk propensity behaviors. The best model showed a significant breakfast-manipulation by fat mass interaction (β = 0.25, SE = 0.10, *t* = 2.40, *P* = 0.019). Specifically, individuals with higher fat mass showed increased risk aversion in high-carb/protein (vs. low-carb/protein) condition, whereas individuals with lower fat mass performed in the opposite way (Fig. 3*A*).

**Fig. 3.**
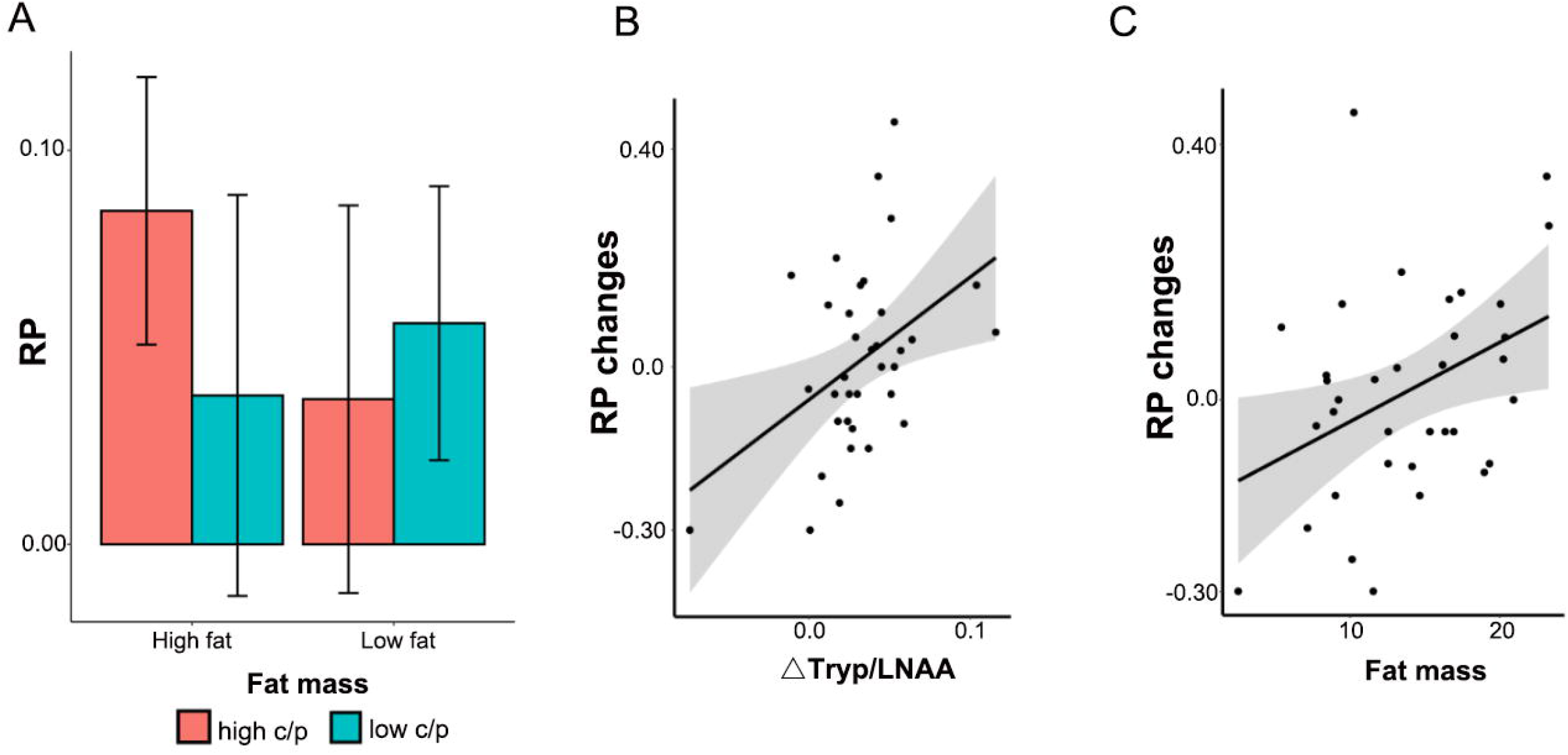
The relationship between tryptophan/LNAA fluctuation, body fat mass and risk propensity changes. (A) The body fat mass modulated the effect of nutrient on risk propensity. Mean and standard error of risk propensities after breakfasts (high and low-carb/protein) in participants with relatively low versus high fat mass on the basis of median split. (B) The tryptophan/LNAA fluctuation affected nutrient-driven changes in risk propensity. (C) The individual fat mass affected nutrient-driven changes in risk propensity. RP, risk propensity.

### The Tryptophan/LNAA affected the Macronutrient Effect on Risky Decision Making

We further identified that 1) the tryptophan/LNAA fluctuation (high-vs. low-carb/protein) predicted the risk propensity changes (*F* = 7.23, *P* = .011, *r^2^* = .18; *r*_spearman_ = 0.37, *P* = 0.030) (Fig. 3*B*); and that 2) the risk propensity changes was also predicted by body fat mass (*F* = 5.77, *P* = .022, *r^2^* = .15; *r*_spearman_ = 0.36, *P* = 0.035) (Fig. 3*C*). Next, we examined the inter-relationship between fat mass, tryptophan/LNAA fluctuation and risk propensity changes by performing a mediation analysis. Table 1 shows that the tryptophan/LNAA fluctuation fully mediated the relationship between body fat and risk-shift. The models with “risk-aversion ➔ tryptophan/LNAA” causal relationship were not considered due to the mismatch to measurement order. Thus, the manipulation of macronutrient exerts its effect on participants’ risk propensity via plasma tryptophan, depending on individuals’ energy storages.

**Table 1.** Mediation analysis: fat mass, tryptophan/LNAA fluctuation, and risk propensity changes.

|  | Path a<br>(X → M) | Path b<br>(M → Y) | Path c<br>(X → Y) | Path c'<br>(X → Y) | Mediation<br>Path (c-c') |
| --- | --- | --- | --- | --- | --- |
| Model 1: X (Fat) → Y (RP) mediated by M (Tryp/LNAA) |  |  |  |  |  |
| $\beta$ | 0.461 | 0.324 | 0.378 | 0.232 | 0.146 |
| $p$ | >0.001 | 0.047 | 0.029 | 0.248 | 0.021* |
| Model 2: X (Tryp/LNAA) → Y (RP) mediated by M (Fat) |  |  |  |  |  |
| $\beta$ | 0.456 | 0.226 | 0.433 | 0.328 | 0.105 |
| $p$ | 0.011 | 0.204 | 0.012 | 0.052 | 0.143 |
| Model 3: X (Tryp/LNAA) → Y (Fat) mediated by M (RP) |  |  |  |  |  |
| $\beta$ | 0.432 | 0.228 | 0.458 | 0.365 | 0.093 |
| $p$ | 0.011 | 0.275 | 0.010 | 0.029 | 0.182 |

RP: risk propensity changes (risk aversion in high-carb/protein vs. low-carb/protein); Tryp/LNAA:
tryptophan/LNAA fluctuation (high-carb/protein vs. low-carb/protein). \* $p < 0.05$ .

### The Macronutrient Effect on Brain Response for Risky Decision Making

We compared the BOLD response during high vs. low risk gamble choices (contrast C2, see *Materials and Methods*) across different sessions. A whole-brain paired t-test revealed increased responses within the right parietal lobule and dorsomedial prefrontal cortex (dmPFC) during the high vs. low-carb/protein sessions (parietal lobule: Montreal Neurological Institute [MNI] coordinates: x = 27, y = −66, z = 42; *t* = 4.24; *P_FDR_* < 0.05; dmPFC: MNI x = 12, y = 54, z = 27; *t* = 3.82; *P_FDR_* < 0.05) (Fig. 4 and *SI Appendix*, Table S4).

**Fig. 4.**
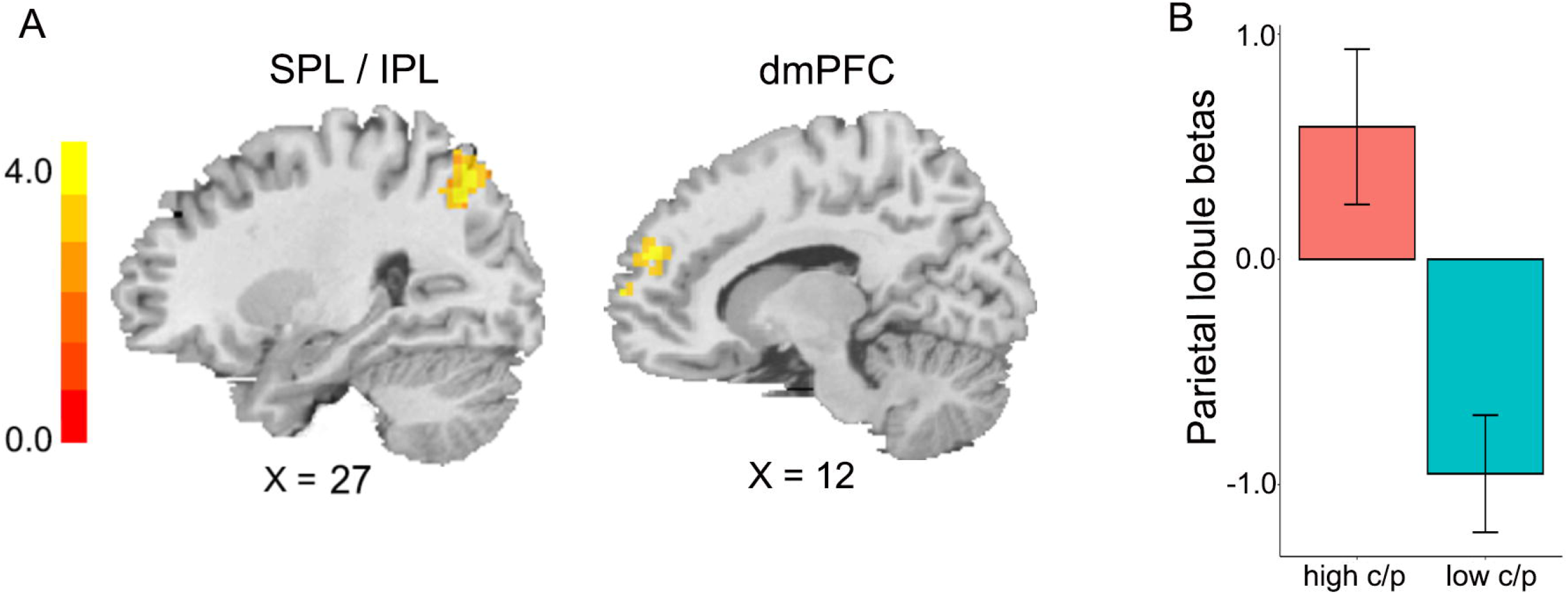
Nutrient modulated brain responses to risk-processing. High-carb/protein (vs. low-carb/protein) breakfast enhanced the parietal lobule activation in risk-processing during gamble choices. (A) Compared to the low-carb/protein session, participants in the high-carb/protein condition showed greater brain responses to risk-processing (contrast C2: high risk > low risk gambles) within the right parietal lobule and dmPFC. (B) Parameter estimates of risk-processing, extracted from the regions of the right parietal lobule showed significant session differences. Error bars are s.e.m. All activations were thresholded using whole brain false discovery rate (FDR) corrected (*P_FDR_* < 0.05). dmPFC, dorsomedial prefrontal cortex; IPL, inferior parietal lobule; SPL, superior parietal lobule.

### The Neural Signal in the Parietal Lobule correlated with Risk Propensity Changes

We tested the relationship between brain activation changes during risk processing (contrast C2: high > low risk gamble) within the right parietal lobule and the dmPFC and the changes of risk propensity between meals (high-vs. low-carb/protein). The result showed that the risk related BOLD changes within the right parietal lobule were significantly correlated with risk propensity changes (parietal lobule: MNI x = 36, y = −75, z = 51, *t* = 3.67; *P_FWE-SVC_* < 0.05) (*SI Appendix*, Table S5).

### The Tryptophan Changes Modulated Neural Signal in the Parietal Lobule

We tested the relationship between macronutrient-affected plasma tryptophan fluctuation and brain activation changes during risk processing (contrast C2: high > low risk gamble) within the right parietal lobule and the dmPFC. The result showed that the high-carb/protein (vs. low-carb/protein) induced tryptophan increase was significantly correlated with risk related BOLD changes within the right dmPFC and parietal lobule (dmPFC: MNI x = 12, y = 66, z = 9, *t* = 3.53; *P_FWE-SVC_* < 0.05; parietal lobule: MNI x =18, y = −63, z = 45, *t* = 3.24; *P_FWE-SVC_ =* 0.07) (*SI Appendix*, Table S5).

A whole-brain regression analysis was also conducted for the relationship between meal-induced tryptophan changes and the brain activation during risk processing. Associations were found between tryptophan fluctuation and risk-related BOLD changes within the bilateral parietal lobule (left parietal lobule: MNI x = −51, y = −42, z = 51, *t* = 5.20; right parietal lobule: MNI x = 30, y = −54, z = 42, *t* = 5.14; *P_FDR_* < 0.05) (Fig. 5*A-B* and *SI Appendix*, Table S6).

**Fig. 5.**
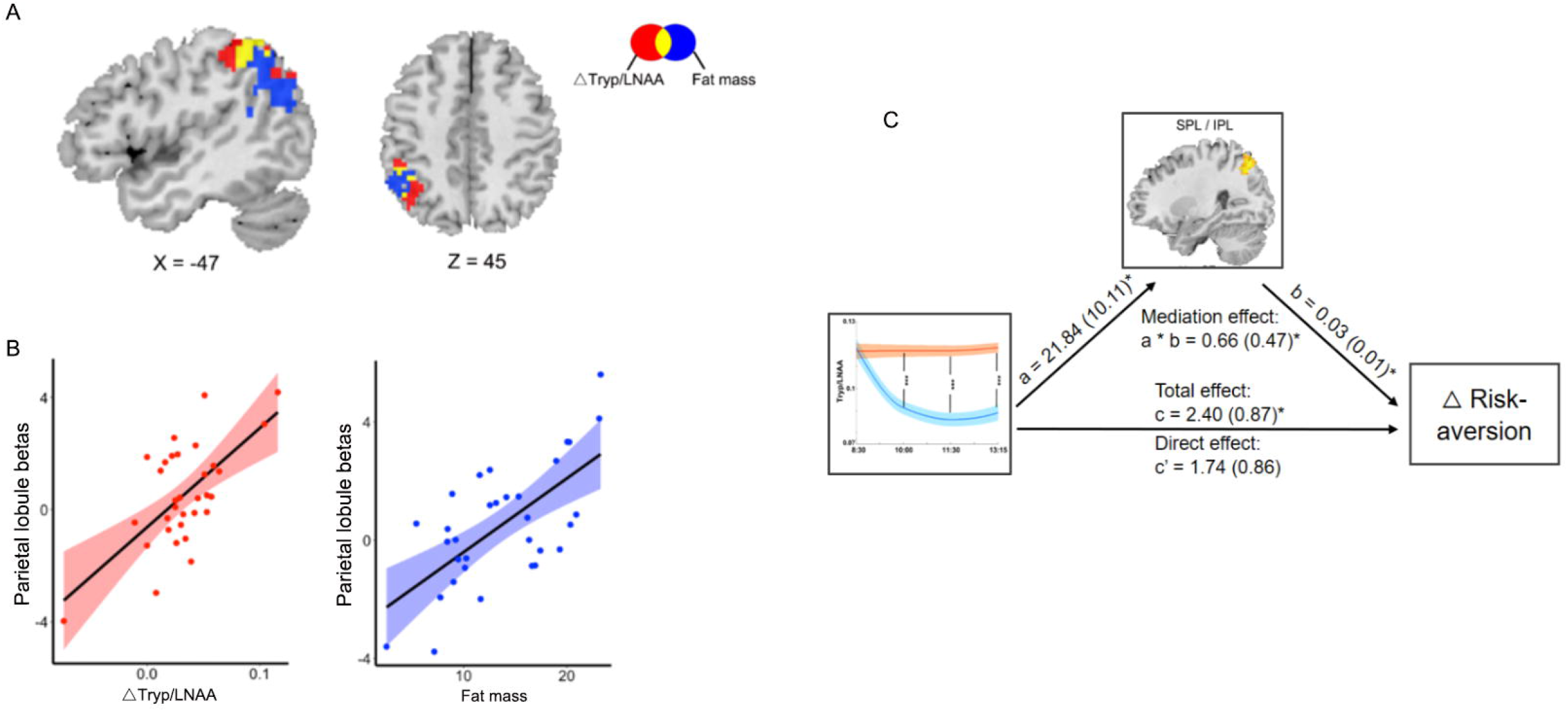
The brain-physiology-behavior relationship between metabolism, body composition and brain response. (A) Positive correlations between tryptophan/LNAA fluctuation and meal-induced BOLD changes during risk processing (contrast C2) in the left parietal lobule (in red); Positive correlations between individuals’ body fat mass and meal-induced BOLD changes during risk processing (contrast C2) in the parietal lobule (in blue); The conjunction area is in yellow. (*P_FDR_* < 0.05). (B) Scatterplots of correlation between parameter estimates of the left parietal lobule and tryptophan/LNAA fluctuation (left), and fat mass (right). (C) Mediation Effect. The parietal lobule fully mediated the influence of nutrient-driven tryptophan/LNAA fluctuation on risk propensity changes. Path coefficients are shown next to arrows with standard errors in parentheses. The direct path c’ is calculated controlling for the mediator. A bootstrapping test was conducted for the statistical significance of the product “a × b”.

### Individual Variation in Fat Mass correlated with Risk Related Activity in the Parietal Lobule

We next tested the relationship between body fat mass and neural responses for risk processing (contrast C2: high > low risk gamble) within the right parietal lobule and the dmPFC, and found no significant correlations within the two ROIs. However, the whole-brain regression analysis identified that individual differences in fat mass were positively correlated with the BOLD changes during risk processing within the left parietal lobule (MNI: x = −39, y = −51, z = 39; *t* = 4.65; *P_FDR_* < 0.05) (Fig. 5*A-B* and *SI Appendix*, Table S6). Furthermore, the risk related brain activity correlated with fat mass showed overlap with that correlated with tryptophan fluctuation within the region of left IPL (cluster size = 85) (Fig. 5*A-B*). Therefore, the parietal lobule may code the links between the individual fat mass, meal-induced plasma tryptophan fluctuation and risk-processing.

### The Parietal Lobule Mediated the Effect of Tryptophan Fluctuation on Risk Propensity changes

The significant associations were observed between the meal affected (high-vs. low-carb/protein) tryptophan fluctuation and the activity of the right parietal lobule during risk-processing, and between the activity of the parietal lobule and the meal affected risk propensity changes. Thus, we conducted a mediation analysis to examine the inter-relationship between tryptophan fluctuation, the parietal lobule activity and risk propensity changes. The result showed that the right parietal lobule β parameter fully mediated the relationship between meal-affected tryptophan fluctuation and risk propensity changes (*a* = 21.75, SE = 10.01, *p* = 0.011; *b* = 0.03, SE = 0.01, *p* = 0.029; *ab* = 0.65, *Z* = 2.07, *p* = 0.038) (Fig. 5*C*). Thus, the high-carb/protein meal modulates risk propensity, via plasma tryptophan fluctuation coded in the parietal lobule.

## Discussion

The modulation of serotonergic system induces risk behavior changes, a mechanism that may depend on energy reserve. Here we utilized nutrition intervention by affecting macronutrient composition of a breakfast to induce changes in brain serotonergic system. Indeed, we observed significant changes on plasma tryptophan/LNAA as metabolic measures, which precisely mapped to the changes in risk propensity. The macronutrient-induced tryptophan/LNAA fluctuation also mediated the relationship between body fat mass and risk propensity changes. Furthermore, our neural results identified the macronutrient-affected BOLD changes in the parietal lobule as a function of risk processing, the activity of which 1) reflected the meal-induced changes in tryptophan/LNAA fluctuation, 2) correlated with individual fat mass, and 3) mediated the relationship between the tryptophan/LNAA fluctuation and risk propensity changes. Also, a similar activation pattern was found in the dmPFC. Taken together, these results deliver a strong evidence of macronutrient-driven changes in modulating metabolism, the brain and behavior, such that it closely mimics pharmacological intervention driven changes.

Our results were consistent with previous studies showing differences in plasma tryptophan and tryptophan/LNAA ratios followed by food consumption in both animals and humans (1, 3). The dietary effects on brain serotonin system depends on 1) the concentration of free plasma tryptophan and 2) the ratio of tryptophan to the other LNAAs sharing the same transport for the blood-brain barrier (31). The high-carbohydrate diet prompt brain uptake of tryptophan by inducing insulin which increases the uptake of other LNAAs into peripheral tissues, thereby decreasing their competition with tryptophan for brain uptake (2). In our study, a significant increase in insulin levels after high-carb/protein meal (vs low-carb/protein) 0.75h after the blood glucose reach peak may indicate the possibility of carb-rich-meal prompted tryptophan brain-uptake via insulin release (see *SI Appendix*). Furthermore, studies investigating the relationship between serotonin and energy homeostasis have reported positive correlations between serotonin levels and obesity, lipid accumulation in animal models (6, 20). Consistently, our study found the macronutrient affected changes in tryptophan/LNAA in relation to body fat mass, suggesting the energy-nutrient interaction in tryptophan metabolism.

Previous studies tackling role of serotonin on decision making, have delivered inconsistent results. Some studies demonstrated that 5-HT enhanced risk-, harm-aversion (5, 13, 32), as well as facilitated behavioral inhibition (10, 11). However, other studies showed no effects or negative effects of serotonin enhancement in risk avoidance performances (14, 15, 33). These inconsistent effects may due to the complexity of serotonergic system itself (9), the efficiency of different manipulation methods for the 5-HT system (e.g., SSRIs versus tryptophan depletion procedure, acute versus chronic manipulation) (34, 35), and the interaction between manipulation methods and individual characteristics (8). For instance, in the study Crockett and colleagues (5) reported that the SSRI’s impact on avoidance behavior is body-weight dependent. Specifically, participants with lower body weight demonstrated risk avoidance due to the higher effective dose of SSRI, whereas participants with higher body-weight became risk prone due to the lower effective dose of SSRI. In line with this, the current study showed that the individual fat mass significantly modulated the nutrient-induced risk propensity changes. Analogously, the lower body fat was linked to less tryptophan fluctuation and mapped to risk prone after high vs. low carb/protein breakfast. Whereas, participants with higher body fat increased risk aversion as predicted by the greater enhancement of tryptophan. Therefore, the body fat may act crucially in tryptophan metabolism and thereby the nutrient-affected risky decisions.

Many previous studies have demonstrated the key role of the parietal lobule and prefrontal cortex (PFC) in risky decision making (26, 36). Consistently, our results found that participants showed greater brain activation when making high vs. low risky options in the parietal lobule and dmPFC driven by nutrient-manipulation. The activity in the parietal lobule also predicted participants’ risk propensity changes. Specifically, the parietal lobule is implicated to modulate the level of uncertainty, which may relate to its role in judgments about probability and derivatives such as reward value and numerical processing (26, 36) Contrary to previous studies (29, 37), the current study, to some extent, isolated the effects of risk-processing from numerical and value computation by conducting a neural contrast between high and low risk gamble trials with equal expected value. Additionally, previous risky decision making studies identified the involvement of the dmPFC, lateral orbitofrontal cortex (lOFC) and inferior frontal gyrus in individual’s risk-processing by comparing participant’s responses to high risky choices compared with low risky or safe choices (25, 28, 38). Further, the activation of the dmPFC and lOFC were modulated by individuals’ risk attitudes (28, 38). Thus, our studies confirmed the crucial role of the front-parietal regions in risk-processing and the sensitivity to macronutrient manipulation.

Furthermore, the front-parietal regions have been implicated with high density of serotonergic receptor neurons (9). For instance, by manipulating the serotonergic system in human, studies revealed activition changes of the PFC, OFC and striatum during decision making (22–24), as well as enhanced activation of the parietal lobule and dlPFC in response for impulsive control (39). Consistently, our results showed that those regions (e.g. the parietal lobule and dmPFC) as a function of tryptophan/LNAA fluctuation mapped to nutrition-affected risk propensity changes. In addition, previous studies suggested associations between body mass and 5-HT receptors density or transporter (SERT) availability in broad cortical regions (i.e., dlPFC, parietal lobule) (40, 41). The front-parietal regions reflecting tryptophan fluctuation and fat mass in our findings supported the findings above, suggesting the interaction of body composition and serotonergic neurotransmission in modulating decision behaviors.

Here, we investigated only normal weight male participants without insulin resistant. However, according to previous studies, individuals with obesity showed greater impulsivity, diminished tryptophan circulation and metabolic response, as well as the increased insulin resistance (42, 43). Therefore, further investigations are required to reveal the underlying mechanisms in diet induced tryptophan metabolism among clinical populations (e.g., obesity and diabetic patients). Furthermore, it is noteworthy, that compared to pharmacological intervention and acute tryptophan load/depletion (5, 23, 24), the nutrient-variation diets exert relatively weaker effects on plasma tryptophan changes, which may lead to the weaker behavioral reflection. Thus, future studies are requested to systematically investigate the relationship between different nutrient-component ratios and decision-making changes.

Overall, the nutrient-variation diets may offer a non-pharmacological approach with high ecological validity for modulating the serotonin function in regulating decision behaviors. Our results deliver a novel perspective for how nutrition can induce brain and behavioral changes, through the dynamic metabolic fluctuation and the energy homeostasis of the organism. This provides wide-ranging implication by offering evidence-based and personalized nutrition recommendation and opening a new avenue for possible treatment strategies for both metabolic and psychiatric clinical population.

## Materials and Methods

### Participants

To investigate the relationship between nutrition intake, body composition and risk decision behaviors, as well as the metabolism and brain mechanisms underlying, we performed a randomized, two-session study in 35 males, normal-weight volunteers (age, 23.82 ± 3.19 years; BMI, 22.96 ± 1.75 kg/m^2^). Every participant underwent a medical screening to exclude metabolic abnormality. The medical screening consisted of a blood sample, a physical examination, and a questionnaire. Exclusion criteria were any abnormalities in the blood results or physical examination, any physical or psychological disease, shifted day/ night rhythm, being a high-performance athlete, BMI under 18 kg/m^2^ or above 30 kg/m^2^, smoking, or food allergies. 32 of these participants (age, 23.85 ± 3.20 years; BMI, 22.85 ± 1.78 kg/m^2^) completed the two sessions of fMRI measurement successfully.

All participants gave written informed consent according to the Declaration of Helsinki. The study was approved by the local Medical Ethical Commission of the University of Lübeck.

### Experimental Design

The study was carried out in a randomized, balanced, within-subject design during two sessions separated by at least 7 days (maximum 9 days). In both sessions, participants received equal-caloric breakfasts (850 kcal). On one day, a high-carb/protein (80% carbohydrates, 10% fats, and 10% proteins) breakfast and on the other day a low-carb/protein (50% carbohydrates, 25% fats, and 25% proteins) breakfast as control was given to the participants. Participants were instructed to consume the whole breakfast (see Fig. 1 and *SI Appendix* for further details).

The study conducted in a research laboratory. At 0745 hours, body composition was assessed by air displacement plethysmography (ADP) using the BOD POD system (BOD POD®, Cosmed, Fridolfing, Germany) (see Methods section in *SI Appendix*).

At 0815 hours, an intravenous catheter was inserted into a vein of the participant’s non-dominant distal forearm and the baseline blood sample was obtained at 0830 hours. At 0845 hours, participants received breakfast according to the high- or low-carb/protein condition respectively. The blood samples drawn at 0830, 0900, 0915, 0930, 1000, 1030, 1130 and 1315 hours (Fig. 1). During the whole procedure subjects could either lie in bed or sit on a chair, but to leave the room or perform any type of physical exercise was not allowed.

At 1145 hours, participants were instructed to complete the risk decision making task (see Fig. 1 and *SI Appendix*) with the fMRI scan. At 1300 hours, participants were guided to a different room where they completed a questionnaire battery including the PANAS (44), the Big Five Inventory (BFI-10) (45), and the behavioral inhibitory system/behavioral approach system (46). These assessments were performed at 1200 hours because the maximum difference in neurotransmitter precursor concentrations between conditions was expected to be present 3.5-4 h after the food intake (1, 4). Metabolic parameters in 1315 (T4) were considered and reported to ensure the nutrition-driven effects lasting until the end of the test battery including the risk task and several questionnaires.

### Blood Samples Acquisition

Twenty-two plasma amino acids as well as leptin and ghrelin were determined from blood samples drawn at 0830, 1000, 1130, and 1315 hours. From the blood samples drawn at 0830, 0900, 0915, 0930, 1000, 1030, 1130, and 1315 hours, the following parameters were determined: glucose, insulin, and cortisol (see *SI Appendix*).

### fMRI Data Acquisition

The imaging was performed on a 3T MRI system (Siemens Magnetom Skyra, Germany) using a 64-channel head-coil. Gradient echo-planar images (EPI) with 58 slices (voxel size: 3 × 3 × 3 mm^3^, field of view: 192 mm, 2000 ms repetition time (TR), 30 ms echo time (TE), no distance factor) were acquired parallel to the commissural line (AC-PC) with the interleaved manner.

### Statistical analyses

The rate of risk choices were analyzed using t-tests and ANOVAs. In each condition, participant’s risk propensity was defined as the proportion of risk-taking trials (29). The identification of four individual risk propensities (two value levels × two risk levels) enabled us to compute the individual risk aversion as the difference between the risk propensities of two decision lotteries with the same expected value (Low risk trial - High risk trial). In other words, more risk prone individuals perceive the increase in risk as less important compared with more risk aversive ones (25).

Generalized mixed effects regression models to postprandial risk propensity levels as the dependent variable were carried out using the lme4 package (https://github.com/lme4/lme4/) in R package. The simplest model given our design was a model containing only treatment as fixed effects regressor (a dummy variable coding for treatment: Treatment, 1, high-carb/protein meal; 0, low-carb/protein meal). The body fat mass and the treatment by fat mass interaction as fixed effects regressors were added to generate complex models. In each model, random effects structure was kept maximal with by subject random intercepts and slopes for treatment (47). Model selection among the four mixed effects models was based on a model comparison approach using the Akaike information criterion (AIC) that estimates the goodness-of-fit of a model based on its likelihood and complexity (*SI Appendix*, Table S3). The best model was a model with the treatment by fat mass interaction effect, as follows:

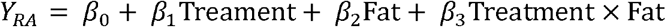

As significant correlations between fat mass, nutrient-driven tryptophan/LNAA fluctuation, and risk-aversion shift during decision-making were observed, we conducted mediation analyses to explore the inter-relationship between these measures using a mediation tool-box (http://wagerlab.colorado.edu/tools) (48). Mediation quantifies the degree in which a relationship between independent variable X and dependent Y, can be significantly explained by a variable M. We defined X as fat mass, Y as risk-aversion shift and M as tryptophan/LNAA fluctuation. A standard mediation model with a bootstrap test (10,000 iterations) for statistical significance was used.

### Metabolic parameters analysis

The influence of breakfast’s macronutrient composition on plasma tryptophan, tyrosine and glucose levels was examined. Ratios between plasma concentrations of tryptophan and tyrosine and the other LNAAs were used as a proxy for brain tryptophan and tyrosine levels and ultimately brain serotonin and dopamine levels (1) (see *SI Appendix*). To examine the effects of nutrient-affected tryptophan on behavioral performance and the brain responses, the tryptophan/LNAA fluctuation ([△tryptophan/LNAA_(T3-T1)_ in the high-carb/protein session] – [△tryptophan/LNAA_(T3-T1)_ in the low-carb/protein]) was used to conduct the Spearman correlation analyses with fat mass and risk propensity changes.

### Imaging data analysis

Spatial processing, subject-level modeling, and group-level statistics were conducted using Statistical Parametric Mapping (SPM12; Functional Imaging Laboratory, London, U.K.). The functional imaging data was spatially realigned, inspected for excessive motion (scans with displacement ≥ 3.0 mm were excluded), normalized to the standard Montreal Neurological Institute EPI template (voxel size = 3 × 3 × 3 mm^3^), and smoothed with an isometric Gaussian kernel (full width at half-maximum of 5 mm). Subject-level models included onsets and durations of events convolved with the hemodynamic response function and motion parameters from realignment, and models were high-pass filtered at 128 s.

Statistical analysis of individual participant imaging data was performed using a general linear model (GLM) of BOLD responses (49). The GLM was estimated to investigate macronutrient modulated risk processing depending on participants’ choice responses. The GLM had 4 regressors: 1) indicator for safe choice trials in low risk condition, 2) indicator for safe choice trials in high risk condition, 3) indicator for gamble choice trials in low risk condition, 4) indicator for gamble choice trials in high risk condition. For this GLM, we calculated the following first-level single-subject contrasts: C1, the overall risk-processing by using the gamble trials minus safe choice trials across risk levels; C2, risk-processing in risk-taking (gamble) trials between high and low risk levels; C3, risk-processing in safe choice trials between high and low risk levels.

A fixed effect model was used to create cross-run contrasts for each subject for a set of contrast images. These contrast images were entered into a second-level random-effects analysis using a paired t-test design to investigate between-condition effects (high-versus low-carb/protein conditions). Whole-brain statistical maps were voxel-level thresholded at *P* < 0.005 prior to undergoing cluster-level false-discovery-rate (FDR) correction (*P_FDR_* < 0.05). The right parietal lobule and dmPFC showed significant session difference in gambling contrast (C2) were then subtracted as masks for region-of-interest (ROI) based regression analyses. The results were visualized using DPABI (http://rfmri.org/dpabi). All reported coordinates (x, y, z) are in Montreal Neurological Institute (MNI) space. See *SI Appendix* for further details.

## Acknowledgements

This work was supported by the German Research Foundation Grants PA 2682/1-1 and PA 2682/2-1 (to S.Q.P.), and the German center for Diabetes Research (DZD) grant 82DZD00902 (to S.M.S.). Both S.Q.P. and L.L. were supported by a grant from the German Ministry of Education and Research (BMBF) and the State of Brandenburg (DZD grant FKZ 82DZD00302).

