## Supplementary material for "Eating to dare - Nutrition impacts human risky decision and related brain function": SI Appendix: Supplemental information.docx

**Supplementary Text**

**Methods**

**Experimental design**

For the two equal-caloric breakfasts, 1) the high-carb breakfast contained: 88 g “Vital-Fit” whole-grain bread, 20 g ham, 5 g cream cheese, 30 g strawberry marmalade, 130 mL milk, 200 mL apple juice, 110 mL water, 225 g banana, and 225 g apple; and 2) the high-protein breakfast contained: 70 g sunflower seed bread, 70 g Vital-Fit whole-grain bread, 40 g ham, 30 g cream cheese, 40 g Camembert, 240 mL milk, 200 mL water, 250 mL yogurt, and 120 g banana.

**Body composition and blood samples measurements**

Body mass was measured directly before subjects entered the chamber using an electronic scale connected to the BOD POD system. While measurements, subjects were sitting in the chamber wearing tight-fitting underwear and a swim cap. Two repeated measurements of body volume were performed. If values deviated more than 150mL, a third body volume measurement was done. Data were averaged and body composition parameters, i.e. body fat mass and fat free mass, were calculated according to manufacturer’s software based on Siri’s equation for body density (version 5.4.1; BOD POD®, Cosmed, Fridolfing, Germany). Thoracic gas volume was predicted. Before each measurement, a two-step calibration was carried out.

All the blood samples were centrifuged at 4 °C and stored at -80 °C until analysis. Blood glucose was measured by an enzymatic-amperometric method (EKF Diagnostic, Barleben, Germany). Insulin and cortisol were assessed by immunoassays (Immulite 2000, Siemens Healthcare Diagnostics, Erlangen, Deutschland). Leptin as well as active ghrelin concentrations were assessed by a radioimmunoassays (RIA Kit, EMD Millipore Corporation, St. Louis, Missouri, USA). Measurement of plasma amino acids was performed according to the method of a previous study (1). Harder and colleagues combine precipitation, derivatization, and chromatographic separation to determine all proteinogenic amino acids, citrulline, and ornithine.

**Risk decision making task**

All participants went through the risk decision making task in the MRI scanner and made repeated choices between a risky option (gamble) and a safe alternative (Fig. 1*B*). The risky options consisted of a 50% gain of either a larger or a smaller option. The safe options consisted of a 100% gain of an intermediate amount. To equate visual stimulation between the safe and the risky options, 2 times the same number was displayed for the safe option. Participants had to indicate their choice by button press during the 4-s presentation. After choice, the chosen option was framed for 0.5 s. No outcome was shown. Intertrial intervals consisted of a fixed part of 2 s and a variable with a high tail and a mean of 2 s (2). At the end of the experiment, one trial was chosen randomly and played out to determine participants’ payoff in Euro. There were four different risky options (gambles) presented (€15/€45, €10/€50, €40/€80, €30/€90), resulting in 2 levels of expected value, 30 and 60€, with the first one offering €15/€45 or €10/€50 gamble and the second one offering €40/€80 or €30/€90 gamble, respectively. There were 20 trials per gamble, with the safe option varied within the range of the risky option it was presented with. Specifically, according to precious studies (3, 4), to ensure that the risky and safe alternatives have approximately the same utility, the safe option was defined by considering the risk variance and individuals’ subjective expected value. The safe alternatives in each gamble, along with their mean, standard deviation and range were: gamble €15/€45: 28 ± 6.37 €, range = 20; €10/€50: 28 ± 9.32 €, range = 30; €40/€80: 56 ± 9.32 €, range = 30; and €30/€90: 56 ± 12.50 €, range = 40. The experiment comprised four sessions each consisting of 20 trials.

**Metabolic parameters analysis**

Ratios between plasma concentrations of tryptophan and tyrosine and the other LNAAs were used as a proxy for brain tryptophan and tyrosine levels and ultimately brain serotonin and dopamine levels. To dynamically track the nutrient-driven metabolic parameters changes, repeated-measures ANOVA was conducted with the time (0830-1315 hours) and session (high-carb/protein, low-carb/protein) as within-subject factors and tryptophan- or tyrosine/LNAA ratios and glucose levels as the dependent variable. Post-hoc tests were performed to investigate the exact relationship between sessions. Next, differences in AUC values (0830-1315 hours) of tryptophan- or tyrosine/LNAA ratios and glucose levels were tested using paired t-tests (two-sided).

**Imaging data analysis**

To probe the links between task performance, tryptophan metabolism, body fat mass and brain responses for risk processing, the individual contrast images for high-carb/protein versus low-carb/protein sessions in gamble trials (C2) were made and voxel-wise regression analyses was conducted within voxels finding session differences in risk-processing. We report signiﬁcant brain activations within the ROIs that survived FWE correction for multiple comparisons using small-volume correction (*P_SVC-FWE_* < 0.05). We also conducted whole-brain regression analyses to assess correlations between brain responses and risk-aversion, tryptophan/LNAA fluctuation as well as fat mass. Moreover, as shown in the Results, we observed significant associations between risk propensity changes, tryptophan fluctuation, and brain activities in the parietal lobule. Thus, a mediation analysis was conducted to explore the inter-relationship between these variables in the mediation tool-box (5). We deﬁned X as tryptophan/LNAA fluctuation, Y as risk propensity changes and M as the activity in the right parietal lobule (mean BOLD parameter estimates at an 8-mm sphere around the peak [36, -75, 51] was extracted).

**Results**

**Metabolic reflection modulated by the breakfasts**

For tyrosine/LNAA, there was a main effect of session [*F*(1, 34) = 7.24, *P* = 0.011] and a trend for an interaction between session and time [*F*(3, 102) = 2.79, *P* = 0.057]. Tyrosine showed a delayed fluctuation, such that its levels were significantly lower in the high-carb/protein condition at T3 [*t*(34) = -3.69, *P* = 0.001] and T4 [*t*(34) = -3.62, *P* = 0.001], but not before [T1: *t*(34) = -0.44, *P* = 0.665; T2: *t*(34) = -0.39, *P* = 0.703] (Fig. S1). For the differences in glucose metabolism between sessions, a paired t test was also conducted to test for changes in glucose increase between 0830 and 0915 hours, which showed a significantly higher peak blood glucose concentrations in the high-carb/protein condition compared with the low-carb/protein breakfast [*t*(34) = 6.31, *P* < 0.001].

We next compared the condition-differences in areas under the curve (AUCs) for glucose, tryptophan and tyrosine levels (T1-T4) using paired t test. We found that, compared with the low-carb/protein breakfast, 1) the high-carb/protein breakfast significantly increased the glucose concentration [*t*(34) = 3.53, *P* = 0.001], 2) it significantly increased plasma tryptophan/LNAA ratio [*t*(34) = 9.06, *P* < 0.001], and 3) significantly decreased plasma tyrosine/LNAA ratio [*t*(34) = -2.66, *P* = 0.012].

No other metabolic parameters [insulin, cortisol, ghrelin, and leptin] were significantly modulated by the different breakfasts (Table S1). However, a significant increase in insulin levels after high-carb/protein meal (vs low-carb/protein) 0.75h after the blood glucose reach peak was found [*t*(30) = 2.05, *P* = 0.049].

For the effect of body composition on tryptophan, a linear regression model was conducted [*Y_Tryp/LNAA_* = *β_0_* + *β_1_Treatment* + *β_2_Fat* + *β_3_Treatment* * *Fat*]. A significant treatment by fat mass interaction confirmed the modulation effect of fat mass on the relationship between macronutrient-manipulation and tryptophan/LNAA fluctuation (β = 0.38, SE = 0.13, *t* = 2.99, *P* = 0.004). Specifically, individuals with higher fat mass showed greater increases in tryptophan/LNAA in high-carb/protein (vs. low-carb/protein) condition (the treatment effect: β = 0.26, SE = 0.05, *t* = 5.20, *P* < 0.001). Whereas, individuals with lower fat mass showed less increases in tryptophan/LNAA fluctuation (β = 0.16, SE = 0.04, *t* = 3.69, *P* = 0.001).

**The influence of macronutrient manipulation on risk behavior**

**Nutrition, body fat and risk propensity**

To test whether and how nutrition intervention and body fat predicts risk-preference, we compared three mixed-effects logistic regression models (Table S3) in their predictive power of the nutrient manipulation on risk behavior. The first model only included fixed effects, and the rest three models included by-subject random intercept and slop. The Akaike Information Criterion (AIC) was used to estimate and compare the goodness-of-ﬁt of the models. Lower AIC values represent a better model fit, while accounting for the number of parameters. According to the AIC values, the best model to predict participant’s risk-behavior was [*Y*_RA_ = *β_0_* + *β_1_Treatment* + *β_2_Fat* + *β_3_Treatment * Fat*], which showed a significant breakfast-intervention by fat mass interaction (β = 0.25, SE = 0.10, *t* = 2.40, *P* = 0.019).

**Fig. S1.** Effects of breakfast-manipulation in tyrosine/LNAA metabolism. Macronutrient composition affected changes in postprandial tyrosine. Red lines indicate high-carb/protein and blue lines indicate low-carb/protein session for tyrosine/LNAA ratio. ^**^*P* < 0.01.

* *

* *

**08:30	10:00	11:30	13:15**

**0.115**

**0.11**

**0.105**

**0.10**

**0.095**

**0.09**

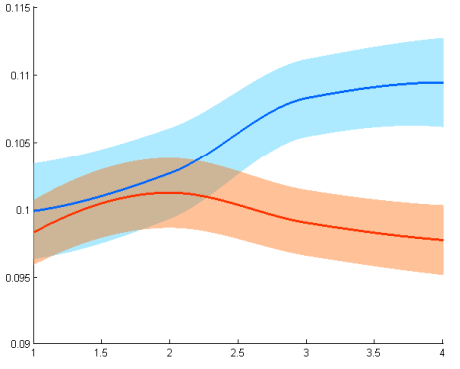

**Table S1** ANOVA results for metabolic parameters.

| **Meals** | **Parameters** | **T1** | **T2** | **T3** | **T4** | **T5** | **T6** | **T7** | **T8** |
| --- | --- | --- | --- | --- | --- | --- | --- | --- | --- |
| High C/P | Cortisol  (Mean±S.D.) | 13.03±3.50 | 12.51±4.49 | 12.84±4.60 | 12.22±4.01 | 11.19±3.94 | 9.62±3.64 | 7.79±2.79 | 10.86±4.00 |
| Low C/P |  | 12.60±2.76 | 12.15±2.84 | 12.45±3.17 | 11.71±3.49 | 10.41±2.89 | 8.58±2.62 | 7.03±2.55 | 9.40±2.92 |
| ANOVA |  | *F*(session) = 2.23 | |  | *F*(time) = 34.42*** | |  | *F*(session*time) = 0.64 | |
| High C/P | Insulin  (Mean±S.D.) | 4.22±1.80 | 13.58±11.73 | 41.55±25.10 | 47.87±28.16 | 43.50±28.76 | 35.66±25.58 | 22.02±18.06 | 6.03±4.31 |
| Low C/P |  | 4.46±1.67 | 13.39±9.17 | 43.15±23.98 | 46.45±27.95 | 34.96±19.49 | 32.25±17.07 | 21.06±15.40 | 7.75±5.30 |
| ANOVA |  | *F*(session) = 0.60 | |  | *F*(time) = 49.92*** | |  | *F*(session*time) = 1.21 | |
| High C/P | Ghrelin  (Mean±S.D.) | 57.36±31.26 |  |  |  | 76.31±37.54 |  | 81.68±44.00 | 128.19±54.39 |
| Low C/P |  | 56.44±21.38 |  |  |  | 89.03±44.57 |  | 65.72±37.17 | 122.04±68.30 |
| ANOVA |  | *F*(session) = 0.41 | |  | *F*(time) = 28.46*** | |  | *F*(session*time) = 1.56 | |
| High C/P | Leptin  (Mean±S.D.) | 6.48±4.02 |  |  |  | 5.67±3.80 |  | 6.05±3.86 | 6.65±4.05 |
| Low C/P |  | 6.74±4.16 |  |  |  | 5.52±3.43 |  | 6.05±3.57 | 6.91±4.61 |
| ANOVA |  | *F*(session) = 0.07 | |  | *F*(time) = 21.15*** | |  | *F*(session*time) = 0.48 | |

T1-T8, the time measurements of blood samples at 0830, 0900, 0915, 0930, 1000, 1030, 1130, and 1315 hours; ghrelin and leptin were determined from blood samples drawn at 0830, 1000, 1130, and 1315 hours (T1, T5, T7 and T8). ^***^*P* < 0.001.

High C/P, high-carb/protein breakfast; Low C/P, low-carb/protein breakfast; SD: standard deviation.

**Table S2** Behavioral measurement: probability of selecting a risky over a safety option in the risk decision-making task.

| **Session** | **Value** | **Low risk**  ***Mean (S.D.)*** | **High risk**  ***Mean (S.D.)*** | **Risk-aversion**  ***Mean (S.D.)*** |
| --- | --- | --- | --- | --- |
| High-c/p | Small EV | 0.51 (0.13) | 0.50 (0.13) | 0.060 (0.178) |
|  | Large EV | 0.51 (0.14) | 0.46 (0.16) |  |
| Low-c/p | Small EV | 0.54 (0.18) | 0.52 (0.15) | 0.047 (0.178) |
|  | Large EV | 0.49 (0.16) | 0.46 (0.14) |  |

EV: expected value; High-c/p: high-carb/protein; Low-c/p: low-carb/protein; SD: standard deviation.

**Table S3** Best mixed effects regression analyses for risk propensity.

| **Models** | **Regressiors** | **Variables** | **AIC** | **LL** | **△AIC** | **ꞷ-AIC** |
| --- | --- | --- | --- | --- | --- | --- |
| 1 | 4 | Treatment, Fat, Treatment × Fat | -44.53 | 27.26 | 11.98 | 0.0016 |
| 2 | 4 | Treatment, Fat, Treatment × Fat^a^ | **-56.51** | **32.26** | **0** | **0.634** |
| 3 | 3 | Treatment, Fat^a^ | -52.88 | 32.44 | 3.63 | 0.103 |
| 4 | 2 | Treatment ^a^ | -54.74 | 32.37 | 1.77 | 0.262 |

Each model contained a fixed-effects intercept in addition to the fixed-effects regressors of interest. In bold is depicted the winning model. AIC, Akaike Information Criterion. LL, loglikelihood estimation.

^a^ Models with by-subject random effects.

**Table S4** Brain regions showing significant differences in response to risk-processing in gamble trials (high > low risk trials) after high- versus low-carb/protein breakfast.

| Regions | Hemisphere | BA | Volume  *(voxel)* | Peak MNI *(mm)* | | | Peak *t* | *P*-value |
| --- | --- | --- | --- | --- | --- | --- | --- | --- |
|  |  |  |  | X | Y | Z | value | cluster(uncorr) |
| **HC > LC** |  |  |  |  |  |  |  |  |
| SPL/ IPL/ SOG/ precuneus/ angular gyrus * | R | 7, 19 | 114 | 27 | -66 | 42 | 4.24 | < 0.001 |
| dmPFC/ SFG/ MFG/ OFC * | R | 10, 9 | 91 | 12 | 54 | 27 | 3.82 | 0.002 |
| fusiform/ parahippocampal gyrus/ ITG | R | 36 | 33 | 39 | -33 | -15 | 5.03 | 0.038 |
| precuneus | R/L | 7 | 31 | 6 | -66 | 42 | 3.73 | 0.043 |

BA: Brodmann’s area; dmPFC: dorsomedial prefrontal cortex; HC: high-carb/protein breakfast; IPL: inferior parietal lobule; ITG: inferior temporal gyrus; L: left; LC: low-carb/protein breakfast; MFG: middle frontal gyrus; OFC: orbital frontal cortex; R: right; SFG: superior frontal gyrus; SOG: superior occipital gyrus; SPL: superior parietal lobule.

* cluster-level false-discovery-rate (FDR) correction (*P_FDR_* < 0.05).

**Table S5** Findings from ROI analyses about regressions between tryptophan/LNAA fluctuation, fat mass and BOLD changes in response to risk-processing in gamble trials (*P_SVC-FWE_* < 0.05).

| **ROI** | **Hemisphere** | ***P_FWE_*** | **Peak MNI *(mm)*** | | | **Peak *t***  **value** |
| --- | --- | --- | --- | --- | --- | --- |
|  |  |  | **X** | **Y** | **Z** |  |
| **Risk propensity changes** |  |  |  |  |  |  |
| parietal lobule | R | 0.028 | 36 | -75 | 51 | 3.67 |
| **Tryptophan/LNAA** |  |  |  |  |  |  |
| dmPFC | R | 0.040 | 12 | 66 | 9 | 3.53 |
| parietal lobule | R | 0.070 | 18 | -63 | 45 | 3.24 |

dmPFC: dorsomedial prefrontal cortex; L: left; MNI: Montreal Neurological Institute space; R: right; ROI: region-of-interest.

**Table S6** Tryptophan/LNAA fluctuation and fat mass modulate BOLD changes in response to risk-processing.

| **Regions** | **Hemisphere** | **BA** | **Volume**  ***(voxel)*** | **Peak MNI *(mm)*** | | | **Peak *t***  **value** | ***P*-value**  **cluster(uncorr)** |
| --- | --- | --- | --- | --- | --- | --- | --- | --- |
|  |  |  |  | ***X*** | ***Y*** | ***Z*** |  |  |
| **Tryptophan/LNAA** |  |  |  |  |  |  |  |  |
| IPL/ angular gyrus/ SMG ^a^ | L | 40,7 | 224 | -51 | -42 | 51 | 5.20 | < 0.001 |
| IPL/angular gyrus/ precuneus/ SPL ^a^ | R | 40, 7 | 130 | 30 | -54 | 42 | 5.14 | < 0.001 |
| MFG/ SFG | R | 9, 8 | 64 | 39 | 33 | 33 | 5.30 | 0.006 |
| MFG | L | 9, 8 | 30 | -30 | 21 | 42 | 4.42 | 0.044 |
| **Body fat mass** |  |  |  |  |  |  |  |  |
| IPL/ angular gyrus/ MTG ^a^ | L | 40,39, 7 | 380 | -39 | -51 | 39 | 4.65 | < 0.001 |
| IFG/ MFG | L | 9 | 74 | -39 | 24 | 30 | 4.34 | 0.004 |
| angular gyrus/ IPL | R | 39 | 42 | 51 | -69 | 39 | 3.65 | 0.021 |
| MFG | L | 8 | 41 | -39 | 18 | 51 | 3.95 | 0.022 |
| IPL/ SMG | R | 40 | 32 | 48 | -42 | 42 | 3.47 | 0.040 |

BA, Brodmann’s area; IFG, inferior frontal gyrus; IPL, inferior parietal lobule; L, left; MFG, middle frontal gyrus; MNI, Montreal Neurological Institute space; MTG, middle temporal gyrus; R, right; SFG, superior frontal gyrus; SMG, supramarginal gyrus; SPL, superior parietal lobule.

^a^ Cluster-level false-discovery-rate (FDR) correction (*P_FDR_* < 0.05).
